# Extracellular vesicles from wild-type Epstein–Barr virus–transformed B-cells export host DNA and EBV EBER1

**DOI:** 10.64898/2026.03.30.715356

**Authors:** Michelle L. Pleet, Rose Peterson, Stephanie Chidester, Emily H. Stack, Madeleine R. Druker, Jubin George, Charlotte V. Dagli, Abaigeal Donaldson, Joanna Palade, Elizabeth Hutchins, Chang-Sook Hong, Nyater Ngouth, Joan Ohayon, Maria Chiara Monaco, Jason Savage, Tapan K. Maity, Weiming Yang, Lisa M. Jenkins, Ru-Ching Hsia, Ionita C. Ghiran, Kendall van Keuren-Jensen, Kory Johnson, Jennifer C. Jones, Steven Jacobson

## Abstract

Epstein–Barr virus (EBV) infection is nearly ubiquitous and strongly linked to multiple sclerosis (MS), but how EBV-infected B cells communicate with distal tissues remains unclear. We performed an integrated multiomic characterization of small extracellular vesicles (sEVs) released from spontaneous lymphoblastoid cell lines (SLCLs) derived from healthy donors and patients with MS, transformed *ex vivo* by endogenous wild-type EBV. Proteomics identified over 6,000 shared proteins enriched in nucleic acid–binding and chromatin-associated factors. EV-associated DNA resolved into two structurally distinct compartments: DNase-sensitive, high–molecular weight DNA associated with the vesicle corona and DNase-resistant, nucleosome-sized (∼130–150 bp) DNA. Both compartments were overwhelmingly host-derived and broadly genomically distributed, whereas EBV DNA was minimal. In contrast, viral RNA cargo was dominated by the EBV noncoding RNA EBER1, which was strikingly enriched across all lines and confirmed within individual vesicles by ddPCR and super-resolution microscopy. EBER1 has previously been detected in MS brain tissue, yet its route to the CNS has remained unexplained. Our findings identify sEVs as a plausible vehicle for disseminating this immunostimulatory viral ncRNA beyond sites of latency, pointing to EV-mediated export of EBER1 as a candidate mechanism linking peripheral EBV infection to distal tissue signaling in MS and beyond.

## INTRODUCTION

Epstein–Barr virus (EBV), a ubiquitous gammaherpesvirus, establishes lifelong latency in memory B-lymphocytes and is etiologically linked to multiple malignancies and autoimmune diseases, including multiple sclerosis (MS).^1–4^ Although epidemiological and longitudinal studies strongly support a role for EBV infection in MS pathogenesis,^5^ the cellular and molecular mechanisms by which EBV-infected B-cells contribute to immune dysregulation remain unclear. One unresolved question is how latently infected B-cells communicate with other immune or tissue-resident cells in the absence of productive viral replication.

Extracellular vesicles (EVs) are nanoscale, membrane-bound particles secreted by all cell types and are increasingly recognized as mediators of intercellular signaling during infection and inflammation.^6,7^ EVs can transport proteins, nucleic acids, and lipids to recipient cells and traverse biological barriers, including the blood–brain barrier (BBB).^8,9^ In viral infections, EVs may contain viral components capable of modulating innate and adaptive immunity independently of infectious virions.^10,11^ However, the integrated viral and host cargo of EVs released from EBV-transformed B-cells has not been comprehensively characterized using unbiased multiomic approaches.

Most studies of EBV biology rely on laboratory-transformed lymphoblastoid cell lines (LCLs) generated using prototype strains such as B95-8, which harbor genomic deletions and do not fully reflect wild-type viral diversity.^12–14^ To better model naturally acquired infection, spontaneous lymphoblastoid cell lines (SLCLs) can be derived *ex vivo* from peripheral blood mononuclear cells through transformation by the donor’s endogenous EBV.^15^ While transcriptomic analyses of such cells have been reported,^16^ the composition of EVs released from these wild-type EBV–transformed B-cells remains unknown.

Importantly, EBV-encoded small RNAs (EBERs), particularly EBER1, have been detected in post-mortem MS brain tissue in prior studies,^17^ suggesting the presence of viral RNA within the CNS, despite the rarity of detectable infectious virus. However, mechanisms by which EBER transcripts might access CNS-resident cells remain unresolved. Because EVs can traverse the BBB and deliver functional nucleic acids to recipient cells, vesicle-mediated transfer represents a plausible pathway linking EBV-infected peripheral B-cells to CNS immune activation.

In the present study, we performed a comprehensive, multiomic characterization of small EVs (sEVs) derived from SLCLs generated from normal donors and patients with stable or active MS. Because EBV is a large DNA virus, we utilized canonical differential ultracentrifugation methods to separate sEVs from virions. By integrating surface profiling, quantitative proteomics, whole-genome sequencing (WGS) of EV-associated DNA, total RNA sequencing (RNASeq), droplet digital PCR (ddPCR), and super-resolution microscopy (dSTORM), we mapped the protein, DNA, and RNA cargo of these vesicles and determined their host and viral origins. Our findings establish a multiomic window into the cargo of EVs from wild-type EBV–transformed B-cells and the data demonstrate export of nucleosome-associated host DNA and the viral noncoding RNA EBER1.

## RESULTS

### Wild-type EBV–transformed B-cells release small extracellular vesicles enriched in B-cell markers

Small EVs were isolated from SLCL culture supernatants by differential ultracentrifugation and analyzed by microfluidic resistive pulse sensing (MRPS), transmission electron microscopy (TEM), and multiplex analysis (MPA) surface profiling (**Fig. 1a**). All SLCLs produced particles with comparable concentrations across donor groups (**Supp. Fig. 1**). TEM revealed characteristic ∼50-200 nm cup-shaped vesicles with similar size distributions, diffuse particulate co-isolates, and no visible intact virions (**Fig. 1b**).

**Figure 1:**
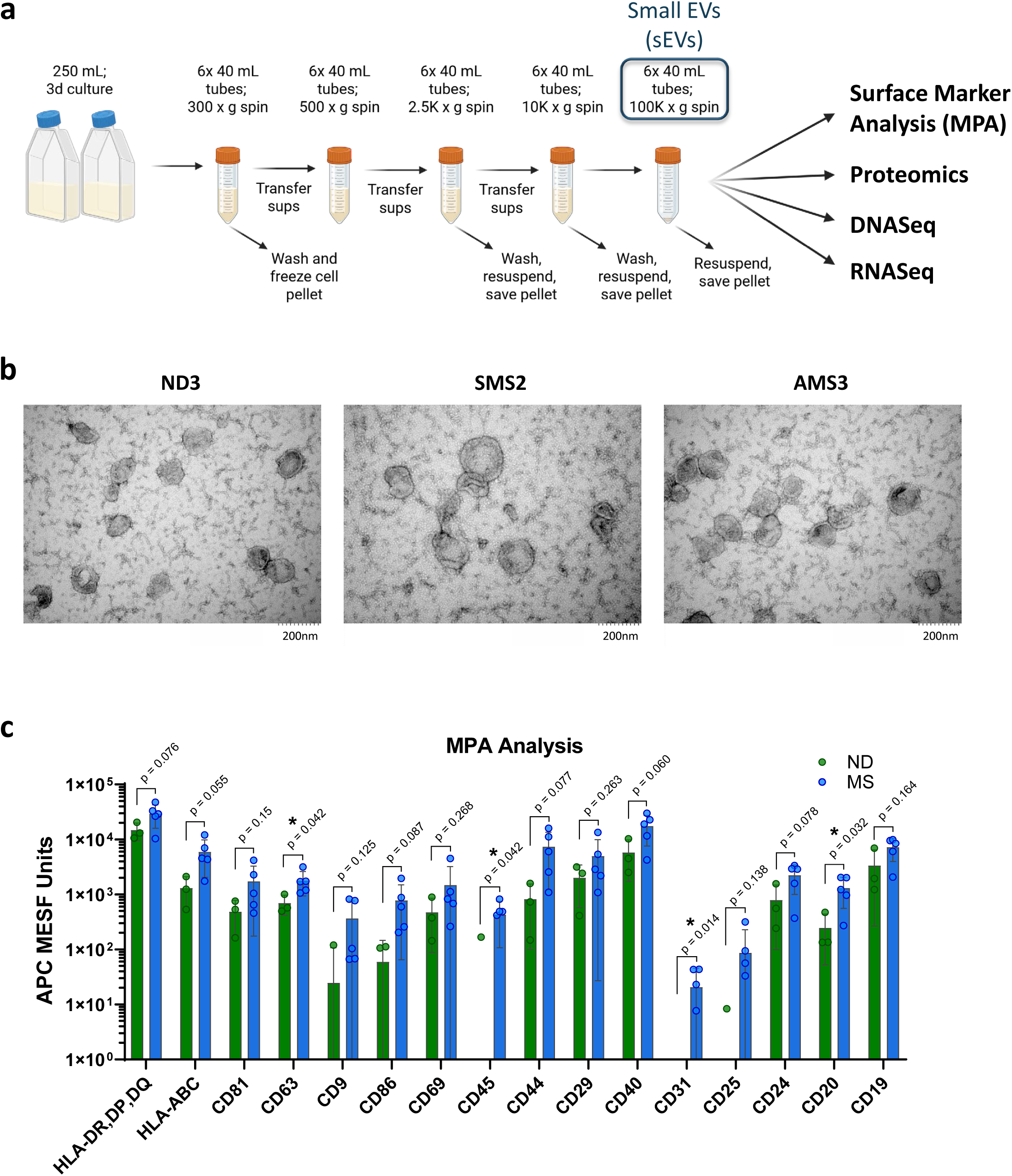
SLCL EV generation, isolation, and phenotyping. **a)** Schematic of the workflow used for small extracellular vesicle (sEV) isolation from spontaneous lymphoblastoid cell line (SLCL) culture supernatants by sequential centrifugation and ultracentrifugation, followed by downstream multiomic characterization including surface marker profiling, proteomics, DNASeq, and RNASeq. Figure was created using BioRender. **b)** Representative transmission electron microscopy (TEM) images of sEV preparations from ND3, SMS2, and AMS3 SLCLs demonstrating vesicles with expected cup-shaped morphology and size range for sEVs (∼50-200 nm in diameter) and no detectable intact virions. Scale bars on the lower right, 200 nm. **c)** Multiplex bead-based surface marker analysis (MPA) of SLCL-derived sEVs. Marker abundance is reported as APC MESF units. P values are shown where indicated, * = p-value ≤ 0.05.

Multiplex bead-based flow cytometry demonstrated robust enrichment of canonical EV tetraspanins (CD9, CD63, CD81) and B-cell markers including MHC-I, MHC-II, CD19, CD20, CD24, and CD40 (**Fig. 1c**). Activation and adhesion molecules (CD44, CD69, CD29, CD45) were also detected. Although some markers were quantitatively increased in MS-derived lines, the overall surface phenotype was consistent across all SLCLs. These findings confirm that spontaneous, wild-type EBV–transformed B-cells release sEVs with preserved B-cell identity.

### Proteomic profiling reveals enrichment of chromatin- and nucleic acid–binding proteins in SLCL sEVs

Data-independent acquisition mass spectrometry (DIA-MS) identified 6,179 proteins shared among SLCL-derived sEVs, establishing a core proteomic signature of EBV-transformed B-cell sEVs (**Fig. 2a**). Principal component analysis (PCA) revealed partial segregation of active MS–derived sEVs from those of normal donors, whereas stable MS samples exhibited greater heterogeneity (**Fig. 2b**).

**Figure 2:**
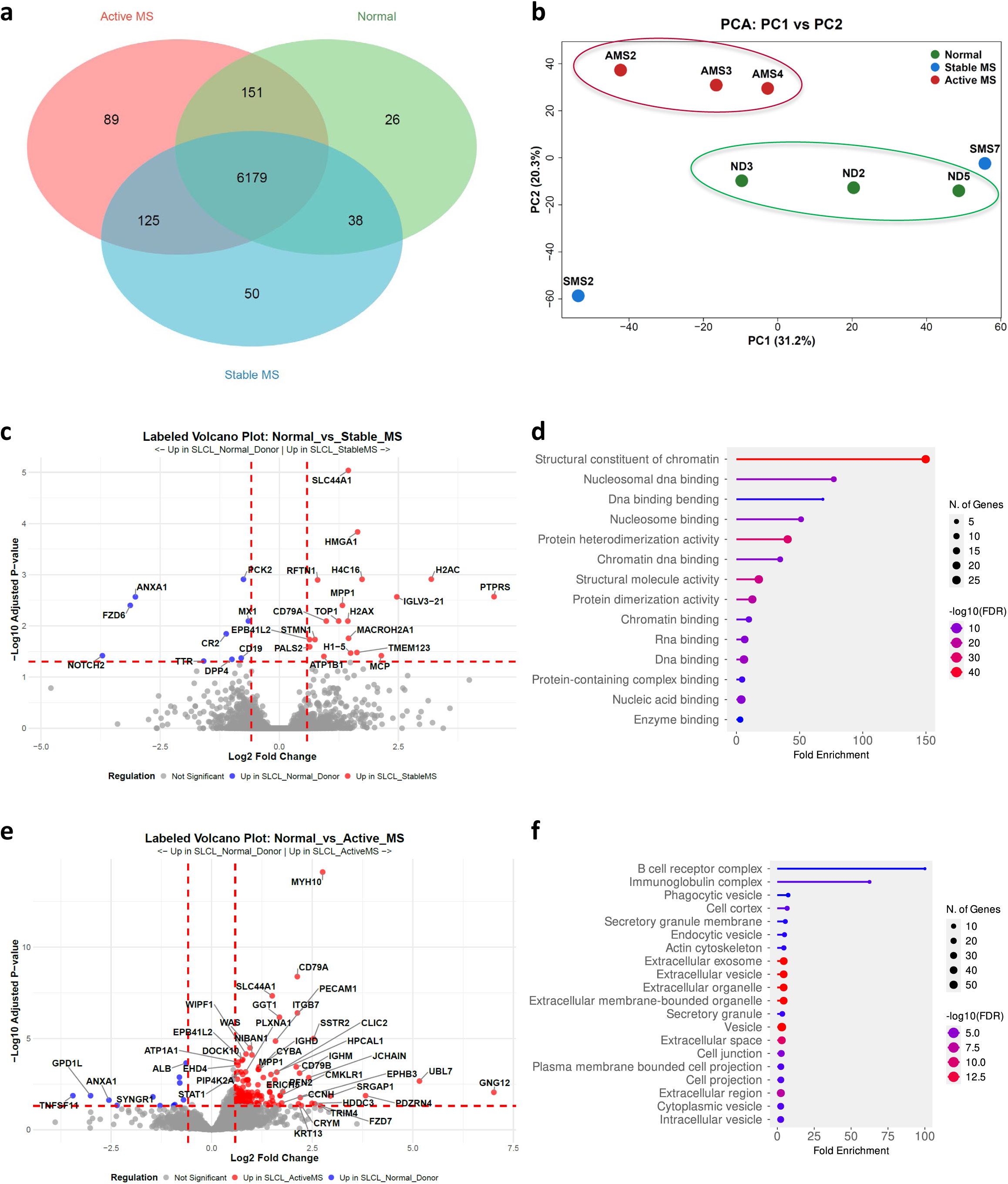
Proteomic profiling of SLCL-derived sEVs. **a)** Venn diagram showing the overlap of proteins detected by DIA mass spectrometry in sEVs derived from ND, SMS, and AMS SLCLs. **b)** Principal component analysis (PCA) of sEV proteomes demonstrating segregation of ND, SMS, and AMS SLCL sEVs based on PC1 (31.2%) and PC2 (20.3%). **c)** Volcano plot comparing SMS to ND sEV proteome differential expression analysis, with top significant differentially abundant proteins labeled. Dotted red lines indicate thresholds chosen for significance (|Log2 fold-change| ≥ log2(1.5); FDR-adjusted p-value < 0.05). **d)** Pathway analysis of significantly increased proteins in SMS-derived sEVs using ShinyGO (version 0.85, https://bioinformatics.sdstate.edu/go/) GO Molecular Function. **e)** Volcano plot comparing AMS to ND sEV proteome differential expression, with top significant differentially abundant proteins labeled. Dotted red lines indicate thresholds chosen for significance (|Log2 fold-change| ≥ log2(1.5); FDR-adjusted p-value < 0.05). **f)** Pathway analysis of significantly increased proteins in sEVs from AMS using ShinyGO (version 0.85, https://bioinformatics.sdstate.edu/go/) GO Cellular Component. Both pathway analysis panels used only the significantly increased proteins with an FDR cutoff of 0.05 and minimum pathway size of 3 proteins.

Differential expression analysis demonstrated that sEVs from stable MS–derived SLCLs were enriched in nucleic acid–binding and chromatin-associated proteins, including multiple histones (H2A, H2AX, H4), topoisomerase 1, and high-mobility group proteins (**Fig. 2c**). Pathway analysis confirmed strong enrichment for nucleosome organization and chromatin-related pathways (**Fig. 2d**). In contrast, sEVs from active MS–derived SLCLs showed enrichment of B-cell receptor components and immunoglobulin-associated proteins (**Fig. 2e, f**).

Notably, 40 EBV-encoded proteins were detected across samples, including EBNA1, LMP2, and major capsid protein (MCP), though viral proteins were not among the most abundant species (**Supp. Table 1**). These data indicate that EBV can drive robust incorporation of host chromatin-associated proteins into EVs and supports the hypothesis that EV cargo may reflect altered nuclear dynamics in transformed B-cells.

### EV-associated DNA exists in two distinct compartments: coronal high–molecular weight DNA and luminal nucleosome-sized DNA

Given the enrichment of chromatin-associated proteins in the sEV proteome (**Fig. 2c, d, Supp. Info. 1**), we next interrogated sEV-associated DNA and distinguished between DNA localized externally on the vesicle surface (coronal) and DNA protected within the vesicle lumen, as these could have different functional effects based on a multitude of factors such as pattern recognition receptors.^18–21^ Analysis of untreated sEV preparations revealed substantial quantities of DNA, including prominent high–molecular weight fragments ranging from approximately 600 to 7,000 base pairs (**Fig. 3a, b**).

**Figure 3.**
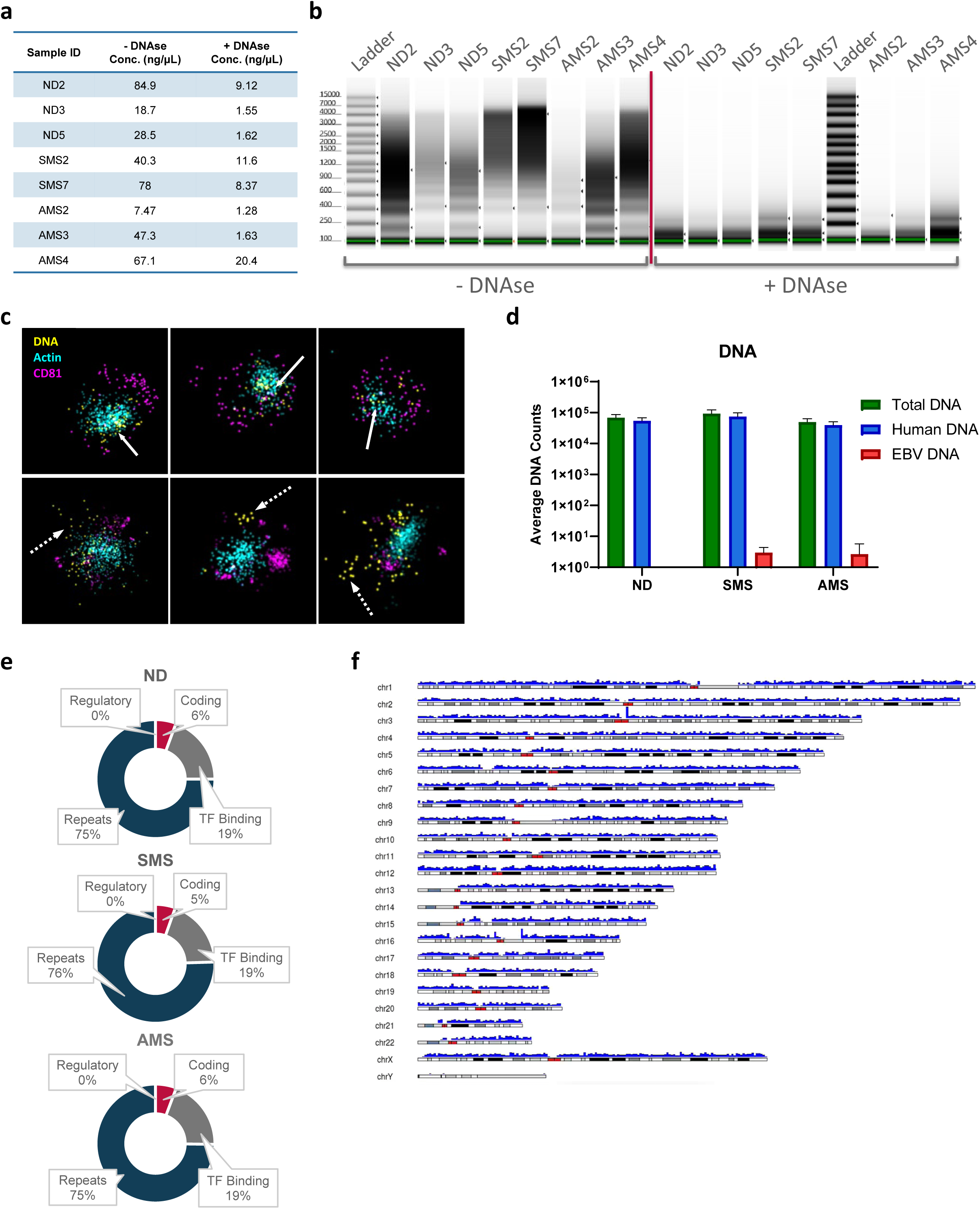
SLCL sEV-associated DNA exists in two distinct compartments and is predominantly host derived. **a)** Quantification of DNA recovered from untreated and DNase-treated sEV preparations from individual SLCL lines by TapeStation gDNA assay, showing substantial reduction in total recoverable DNA after DNase treatment. **b)** TapeStation profiles of sEV-associated DNA before and after DNase treatment. Untreated samples contain abundant high-molecular-weight DNA, whereas DNase treatment removes the majority of this externally accessible fraction and reveals a persistent DNase-resistant population centered at approximately 130-150 bp, consistent with nucleosome-sized luminal DNA. **c)** Representative ONi super-resolution dSTORM images of SLCL-derived sEVs captured by tetraspanin markers and stained for EdU-labeled DNA (yellow), actin (blue), and CD81 (magenta), demonstrating both protected/luminal (solid white arrows) and vesicle surface-associated (dashed arrows) EdU-positive DNA. **d)** Quantification of total, human, and EBV-mapped DNA sequencing reads in averaged ND, SMS, and AMS sEV samples ± SD, showing that EV-associated DNA is overwhelmingly host derived, with minimal EBV genomic DNA. **e)** Donut plots summarizing genomic annotation categories of host-derived SLCL sEV DNA reads for ND, SMS, and AMS groups, demonstrating predominant representation of repetitive elements with smaller contributions from transcription factor (TF)-binding regions, coding regions, and regulatory elements. **f)** Representative karyotype density plot showing genome-wide distribution of host-derived SLCL sEV DNA reads (blue vertical bars) across human chromosomes. Red blocks show centromeric regions of each chromosome. Black and light-gray blocks represent the chromosome ideogram/cytoband structure.

To distinguish coronal (surface-associated) from luminal (intravesicular) DNA, intact sEVs were treated with DNase prior to DNA extraction. DNase treatment markedly reduced the high–molecular weight DNA population, indicating that these fragments were externally accessible and associated with the vesicle corona (**Fig. 3a, b**). Importantly, a distinct population of DNA fragments centered at ∼130–150 base pairs persisted following DNase treatment. This fragment size corresponds closely to DNA protected within nucleosomes, suggesting that chromatinized DNA is present within the lumen of intact sEVs and is shielded from enzymatic digestion. Quantitative assessment demonstrated that both coronal and luminal DNA were present at substantial levels across all SLCL lines, irrespective of donor disease status (**Fig. 3a, b**). Single vesicle, super-resolution microscopy (**Fig. 3c**) was performed to evaluate DNA localization for sEVs. We observed DNA present both on the surface (in proximity with CD81, **Fig. 3c**, dashed arrows) and protected within the sEV lumen (in proximity with actin, **Fig. 3c**, solid arrows). Thus, DNA in SLCL sEVs is present both as luminal fragments and high–molecular weight DNA associated with the vesicle surface.

WGS of sEV-associated DNA revealed that the DNA was overwhelmingly host-derived (**Fig. 3d**). Reads mapped broadly across all chromosomes without preferential enrichment of specific loci and were enriched for repetitive elements (**Fig. 3e, f**). Only minimal EBV genomic DNA was detected, and no consistent enrichment of viral genomic regions was observed (**Fig. 3d, Supp. Fig. 2**). Together, these data demonstrate that EVs released from wild-type EBV–transformed B-cells carry two spatially and structurally distinct pools of host-derived DNA: (i) high–molecular weight DNA associated with the vesicle corona and (ii) nucleosome-sized DNA protected within the vesicle lumen.

### RNA sequencing identifies diverse viral transcripts with striking enrichment of EBER1

Total stranded RNASeq identified abundant host and EBV-derived transcripts within SLCL-derived sEVs (**Fig. 4a**). Protein-coding mRNA comprised the majority of host RNA species across all groups, although diverse noncoding RNA species were also detected (**Fig. 4b**). Chromosomal mapping revealed broad distribution of host RNAs without strong positional bias (representative karyotype density plot in **Supp. Fig. 3**).

**Figure 4:**
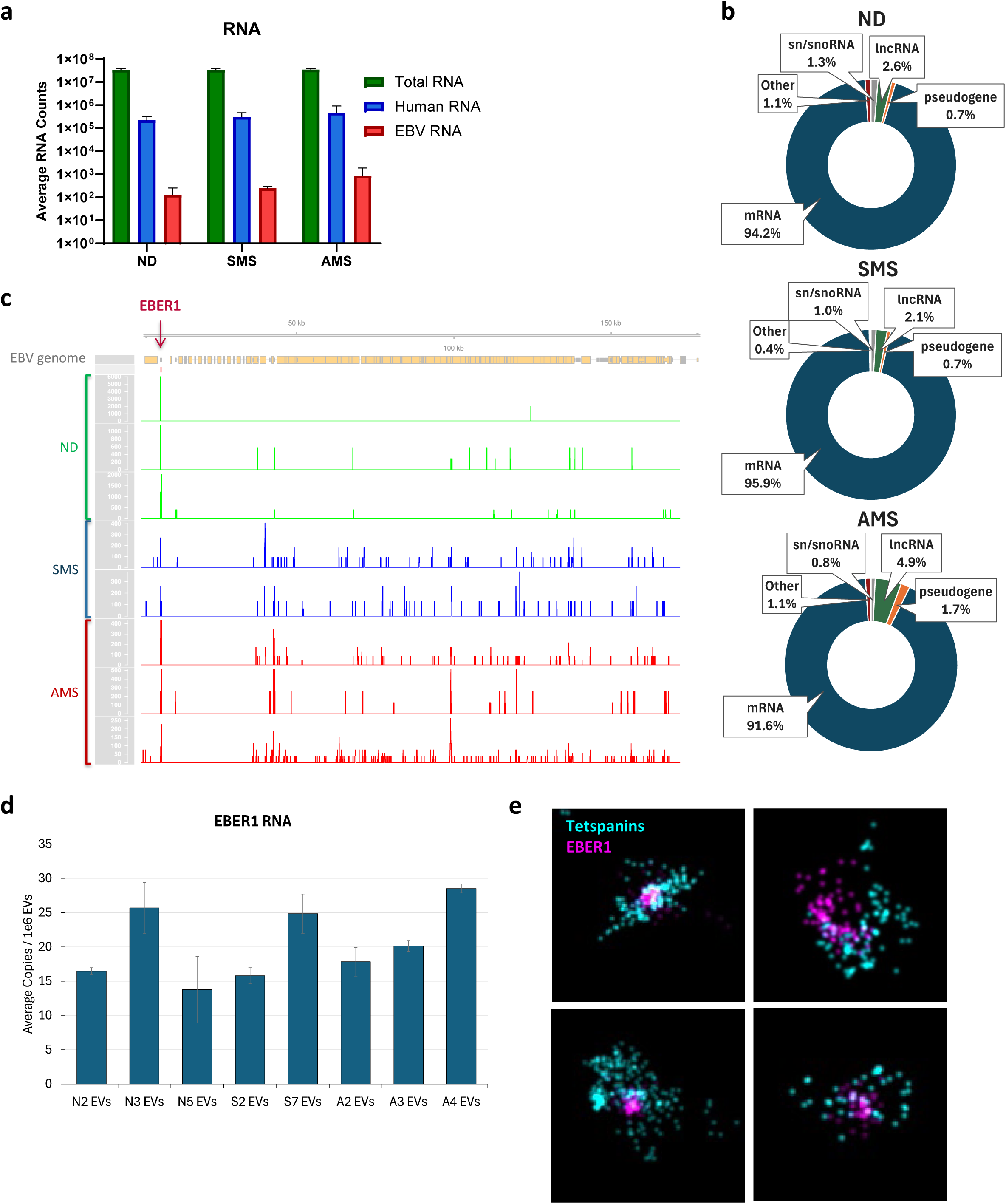
RNA sequencing identifies diverse host and EBV transcripts in SLCL-derived sEVs, with marked enrichment of EBER1. **a)** Quantification of total, human, and EBV RNA reads detected from averaged ND, SMS, and AMS sEVs ± SD, demonstrating abundant host RNA with readily detectable EBV-derived transcripts. **b)** Donut plots showing host RNA biotype composition in ND, SMS, and AMS sEVs. Protein-coding transcripts (mRNA) comprise the majority of mapped host RNA, with smaller contributions from noncoding RNA (ncRNA), small nuclear/small nucleolar RNA (sn/snoRNA), pseudogene transcripts, and other RNA classes. **c)** Integrated Genome Viewer (IGV) browser view of EBV genome-aligned RNASeq reads from individual sEV preparations, illustrating the distribution of viral transcripts across the EBV genome and highlighting strong enrichment of EBER1 (red arrow). **d)** Droplet digital PCR quantification of EBER1 from sEV RNA from individual SLCL lines, reported as average copies per 10^6^ EVs input. **e)** Representative ONi super-resolution microscopy images of SLCL sEVs stained for EBER1 (magenta) and tetraspanins (blue), confirming incorporation of EBER1 ncRNA within vesicles.

Among viral transcripts, 59 EBV-encoded RNAs were detected (**Fig. 4c, Supp. Table 2**). Of note, the viral ncRNA EBER1 was the most abundant EBV transcript in EVs across all donor groups (**Fig. 4c, red arrow**). EBV transcript abundance was increased in MS-derived samples, particularly in active MS, although the qualitative viral RNA profile was shared. Droplet digital PCR confirmed the presence of EBER1 at 14–29 copies per 10^6^ vesicles (**Fig. 4d**), whereas lytic transcripts (e.g., BZLF1/Zta, BFRF3) were detected at less than one copy per 10^6^ vesicles (**Supp. Fig. 4**). Super-resolution microscopy further demonstrated EBER1 localization within individual vesicles, where membranes are identified with a cocktail of antibodies against the tetraspanins CD9, CD81, and CD63 (**Fig. 4e**).

Together, these results indicate that EBV-transformed B-cells selectively enrich EBER1 within EVs while exporting minimal viral genomic DNA. Given prior reports of EBER1 detection in MS brain tissue, the robust incorporation of EBER1 into SLCL-derived EVs suggests a potential mechanism by which EBV-infected B-cells could disseminate viral ncRNA beyond latency sites.

## DISCUSSION

This study provides the first integrated multiomic characterization of extracellular vesicles released from B-cells transformed by endogenous, wild-type EBV by profiling surface markers, proteome, DNA, and RNA cargo. A central finding of this study is that sEV-associated DNA in wild-type EBV–transformed B-cells exists in two discrete compartments: an external, high–molecular weight coronal fraction and an internal, nucleosome-sized luminal fraction (**Fig. 3**). This distinction is mechanistically important. The high–molecular weight DNA that is DNase-sensitive appears to represent co-isolated DNA fragments as well as DNA physically associated with the vesicle surface. EV coronas are increasingly recognized as dynamic molecular interfaces capable of modulating vesicle biodistribution and receptor engagement. The presence of abundant, genome-derived DNA in the vesicle corona suggests that externalized chromatin fragments could directly engage extracellular DNA sensors or cell-surface pattern recognition receptors prior to vesicle internalization. In contrast, the DNase-resistant ∼130–150 bp DNA fragments are consistent with nucleosome-protected DNA packaged within the vesicle lumen. The concomitant enrichment of histones and chromatin-associated proteins in the EV proteome (**Fig. 2c, d**) strongly supports the interpretation that these luminal DNA fragments are associated with nucleosomal complexes. The export of chromatinized DNA within vesicles may potentially suggest a regulated process linked to DNA repair in the donor cells.

Strikingly, both coronal and luminal DNA pools were overwhelmingly host-derived, whereas viral genomic DNA was minimally represented (**Fig. 3d**). Despite the presence of EBV episomes within SLCL nuclei, viral genomic material was not preferentially incorporated into sEVs. The coexistence of surface-associated genomic DNA and luminal nucleosome-sized DNA has important implications for EV biology. Coronal DNA may influence vesicle stability, cellular uptake, and immune recognition extracellularly, whereas luminal chromatin fragments may engage intracellular DNA-sensing pathways following vesicle internalization. Thus, EV-associated DNA may function in both extracellular and intracellular signaling contexts. Whether these DNA populations arise from regulated chromatin remodeling, nuclear blebbing, or vesicle–nucleus crosstalk remains to be determined. This dual localization suggests multiple potential mechanisms by which EVs could influence recipient cells.

In contrast to the minimal representation of viral genomic DNA compared to host DNA, viral RNA was readily detectable (**Fig. 4a**). Notably, the EBV ncRNA EBER1 was consistently the most abundant viral RNA identified across SLCL sEVs (**Fig. 4c**). EBER1 is among the most highly expressed EBV transcripts^22,23^ and has been implicated in innate immune activation through pattern recognition receptors RIG-I and TLR3.^24,25^ Its robust incorporation into extracellular vesicles raises the possibility that EBV-infected B-cells disseminate viral ncRNA through vesicle-mediated transfer, independent of infectious virions. Super-resolution imaging and quantitative ddPCR (**Fig. 4d, e**) confirm that EBER1 is present within individual vesicles. This selective packaging of EBER1 in EVs may not only have broad implications for EBV-related immune responses but also suggests a potential mechanism whereby EBV-infected cells may transfer bioactive cargo to non-immune cells. Given that EVs can traverse biological barriers including the BBB, these vesicles may represent a previously underappreciated route of communication between EBV-infected B-cells and distant immune or neural cells.

The enrichment of EBER1 within EVs is particularly notable in the context of prior reports detecting EBER transcripts in MS brain tissue.^17,26,27^ As productive EBV infection is rarely and controversially observed within the CNS,^28,29^ the presence of viral RNA has remained mechanistically unexplained. Our findings raise the possibility that EBER1 may be delivered to CNS-resident cells through vesicle-mediated transfer from peripherally infected B-cells. EVs are readily capable of crossing the BBB,^30,31^ and EV-associated RNAs can remain functional in recipient cells.^32,33^ The structural incorporation of EBER1 within EVs provides a biologically plausible mechanism linking peripheral EBV latency to viral RNA detection within the MS brain.

Several limitations warrant consideration. The number of donor-derived lines was modest, and differential ultracentrifugation cannot fully exclude all non-vesicular complexes. Functional consequences of EV-associated DNA and EBER1 were not directly addressed here. Nonetheless, the comprehensive multiomic information presented provides a deep catalog of vesicle cargo and identifies avenues for future mechanistic studies. Our study also has the strength of examining EVs from cells infected with wild-type, naturally acquired EBV rather than laboratory strains. Prior studies of EVs from EBV-transformed LCLs generated with laboratory strains have reported selected viral cargoes, including LMP1, EBV miRNAs, EBERs, and some latent mRNAs,^32,34–36^ whereas the sEVs from SLCLs in our study were associated with a broader repertoire of viral RNAs and proteins. This difference may reflect, at least in part, that our RNAseq library approach under-represents small RNAs, as well as the fact that the commonly used laboratory strain B95-8 is a deletion derivative that lacks an approximately 12-kb region containing most BART miRNAs and the LF1, LF2, and LF3 open reading frames.^37,38^ Taken together, our results suggest that wild-type EBV reshapes EV cargo in many ways that may influence intercellular communication beyond the infected cell.

In summary, we define the multiomic landscape of EVs released from wild-type EBV–transformed B-cells and reveal that these vesicles carry structurally distinct pools of host DNA and are selectively enriched in the viral ncRNA EBER1, while containing minimal viral genomic DNA. By identifying EVs as a vehicle for EBER1 export, our findings offer a mechanistic framework through which EBV-infected B-cells may disseminate viral RNA to distal tissues, including the CNS, in the absence of infectious virus. These results provide a resource for understanding how EBV may reshape intercellular communication and contribute to immune-mediated disease.

## MATERIALS AND METHODS

### Cell Culture and Extracellular Vesicle Isolation

Spontaneous lymphoblastoid cell lines (SLCLs) were generated as previously described.^15,16^ Cells were maintained in RPMI-1640 supplemented with 10% fetal bovine serum (FBS), 1% penicillin/streptomycin, and 1% L-glutamine at 0.5–1.5 × 10 cells/mL. For cells grown in EdU (5-ethynyl-2’-deoxyuridine, a nucleoside analog of thymidine) to label newly synthesized DNA, the Click-iT Plus EdU Cell Proliferation Kit (ThermoFisher; Alexa Fluor 488 dye) was used according to manufacturer’s instructions. Briefly, EdU was added to the cell culture media at a final concentration of 10 µM and cells were cultured and fed according to normal conditions for 5-8 days until supernatant/EV harvest.

For EV production, cells were washed and plated at 5–8 × 10 cells/mL in serum-free, phenol red–free XVIVO-15 medium supplemented with 1% penicillin/streptomycin and 1% L-glutamine. After 72 h, conditioned media (∼250 mL) were collected and subjected to differential ultracentrifugation. Supernatants were sequentially centrifuged at 300 × g (10 min), 500 × g (10 min), 2,500 × g (20 min, 4°C), and 10,000 × g (20 min, 4°C). Pellets from the 2.5K and 10K spins were washed once in 0.1 µm–filtered PBS and stored. The final supernatant was ultracentrifuged at 100,000 × g for 2 h at 4°C. The 100K pellet (small EV fraction) was washed in 0.1 µm–filtered PBS, recentrifuged at 100,000 × g for 70 min, and resuspended in filtered PBS for downstream analyses.

### Microfluidic Resistive Pulse Sensing (MRPS)

EV size distribution and concentration were measured using a Spectradyne ARC instrument with TS-400 cartridges. Samples were spiked with 240 nm NIST-traceable beads (1 × 10 particles/mL final concentration) for inter-run normalization. Data were processed in MATLAB (v2023a), and particle concentration was normalized to bead recovery. Statistical analyses were performed in GraphPad Prism (v10.3.1).

### Multiplex EV Surface Profiling

Surface marker analysis was performed using the MACSPlex Exosome Kit (Miltenyi Biotec). EV input (1 × 10□ or 1 × 10□ particles, quantified by MRPS) was incubated overnight with capture beads at room temperature under rotation. Beads were washed and stained with APC-conjugated anti-CD9, anti-CD63, and anti-CD81 detection antibodies according to the manufacturer’s protocol. Samples were acquired on a Cytek Aurora spectral flow cytometer. Instrument calibration and compensation were performed using manufacturer-provided controls and single-stain beads. Data were analyzed using FlowJo (v10).

### Transmission Electron Microscopy

EV morphology was assessed by transmission electron microscopy (TEM). EVs were prepared for transmission electron microscopy by negative staining on glow-discharged, carbon-coated 400-mesh copper grids. Briefly, 5 µL of sample was applied to the grid and allowed to adsorb for 30–60 s. Excess liquid was removed with filter paper, and grids were optionally washed twice with water. Grids were then stained with a filtered negative stain by sequential application to fresh droplets, with a total staining time of approximately 1 min. Excess stain was removed by wicking from the grid edge, and grids were air-dried with the sample side facing upward before imaging. Freshly prepared or clarified stain was used to minimize precipitate formation. Representative images were acquired at multiple magnifications.

### Mass Spectrometry

Isolated EVs were lysed in 5% SDS and digested with trypsin using DNA mini-prep columns, as previously described.^39^ Briefly, samples were reduced with 5 mM Tris(2-carboxyethyl)phosphine (TCEP) and alkylated with 20 mM iodoacetamide. Proteins were then acidified with 2.5% phosphoric acid, diluted in 100 mM triethylammonium bicarbonate (TEAB) (pH 7.55) in 90% methanol, loaded onto columns, washed, and digested overnight at 37 °C with trypsin/LysC. Peptides were sequentially eluted from the S-trap with 50 mM TEAB (pH 8.5), 0.2% formic acid, and 50% acetonitrile in water, then pooled and lyophilized.

Mass spectrometry analysis was performed using a Vanquish Neo UHPLC system (ThermoFisher Scientific) coupled to an Orbitrap Exploris 480 mass spectrometer (ThermoFisher Scientific) operating in data-independent acquisition (DIA) mode. Peptides were separated on a 75 μm × 50 cm EASY-Spray™ Neo UHPLC column (ThermoFisher Scientific). The mobile phases consisted of (A) 0.1% formic acid (FA); and (B) 0.1% FA, 80% acetonitrile. Peptides were separated using a 130 min gradient at a constant flow rate of 300 nl/min: 2% B for 1 min, 2–32% B for 121 min, 32–90% B for 20 min. The Orbitrap Exploris 480 mass spectrometer was operated in positive ion mode with full-scan precursor spectra (MS1) acquired at a resolution of 120,000 over the m/z scan range of 420–680. The AGC target was set to 300% (3e6 ions absolute) with a maximum injection time of 100 ms. Fragment ion scans (DIA MS2) were collected using 60 isolation windows with a 4 m/z window width, covering a precursor range of 430–670 m/z. DIA windows were placed using the instrument’s automatic window-placement optimization. Fragment spectra were recorded in the Orbitrap at a resolution of 90,000, using stepped higher-energy collisional dissociation (HCD) with 25%, 27.5%, and 30% normalized collision energies. The DIA AGC target was set to 3000% (3e6 ions absolute) and the injection time was automatically determined. The scan range for fragment ions was 200–1800 m/z with DIA cycle time of 3 seconds.

DIA raw files were analyzed using DIA-NN 2.2.0^40^ using an in-silico DIA-NN predicted spectral library from the UniProt Homo sapiens and EBV databases, allowing for cysteine carbamidomethylation and N-terminal methionine excision and 1 missed cleavage. The DIA-NN search included the following settings: Protein inference□ = □Genes, Neural network classifier□ = □Single-pass mode, Quantification strategy□ = Robust LC (high precision), Cross-run normalization□ = □RT-dependent, Library Generation□ = □IDs, RT and IM Profiling and Speed and RAM usage□ = □Optimal results. Mass accuracy and MS1 accuracy were set to 0 for automatic inference. No share spectra, Heuristic protein inference and MBR were checked.

### Proteomics Analysis Pipeline

Proteomics analysis was performed in R (v4.5.1) with RStudio (v2024.12.0.467) using the pipeline shown in **Supplemental Figure 5a**. Undetected values were imputed with the placeholder value of 100, and proteins with values greater than 100 were considered detected. Detected values were log2-transformed for downstream analyses. Initial quality control was performed using raw log2-transformed intensities, and sample-level summary statistics, including median and mean log2 intensity and the number of detected proteins per sample, were calculated using detected values only. Protein detection performance was then assessed within each cell line by calculating the proportion of samples in which each protein was detected, and proteins were retained for downstream analysis if they were detected in at least 50% of the 6 replicates within at least one cell line. Quality control visualizations included intensity distributions, detection summaries, sample correlation heatmaps, and Venn diagrams of proteins detected across SLCL donor groups (**Fig. 2a**). PCA was performed on proteins meeting this detection threshold.

For advanced filtering and differential analysis, proteins were further retained if they had at least 3 detected values across the 6 replicates in at least one cell line, and expression matrices were analyzed using the limma/voom workflow to model the mean-variance relationships and assign precision weights for differential abundance testing. Additional PCA and one-way ANOVA analyses were performed on the filtered dataset to evaluate sample structure and exploratory protein-level differences among donor groups. Differential protein abundance analysis was performed using limma (v3.64.3) with empirical Bayes moderation and duplicateCorrelation to account for non-independence among technical replicates from the same cell line. Protein-level expression matrices were generated from the filtered log2-transformed abundance data, and proteins with fewer than 4 non-missing observations in a given comparison were excluded. A no-intercept design matrix was constructed using donor group labels, and contrasts were fit for “Normal vs. Stable MS” and “Normal vs. Active MS” comparisons. Statistical significance was defined as an adjusted P value < 0.05 and an absolute log2 fold change greater than log2(1.5). Significant proteins from Stable MS and Active MS comparisons were imported into ShinyGO (v0.85; https://bioinformatics.sdstate.edu/go/) for GO Cellular Component and GO Molecular Function pathway enrichment analysis. Results were generated in R using tidyverse (v2.0.0), readxl (v1.4.5), and edgeR (v4.6.3). Custom R scripts for proteomics analysis were developed with assistance from Claude (claude.ai; Anthropic; version Claude Sonnet 4.5). The authors verified the code, executed all analyses, and confirmed the results.

### DNase Treatment and EV-Associated DNA Extraction

Intact EV preparations were treated with Heat&Run DNase (ArcticZymes) for 20 min at 37°C followed by enzyme inactivation at 58°C for 5 min. DNA was extracted immediately using the Plasma/Serum Cell-Free Circulating and Viral Nucleic Acid Purification Kit (Norgen Biotek). DNA was quantified by PicoGreen fluorescence assay and fragment size was assessed using Agilent TapeStation (Genomic DNA and D5000 High Sensitivity assays).

### DNA Library Preparation and Whole-Genome Sequencing

EV DNA was isolated from extracellular vesicles using the Plasma/Serum Cell-Free Circulating and Viral Nucleic Acid Purification Mini Kit (Norgen Biotek). Prior to column purification, lysates were treated with RNase A (ThermoFisher Scientific). DNA quality and size distribution were evaluated using TapeStation Genomic DNA and High Sensitivity D5000 assays (Agilent Technologies).

DNA was fragmented to 200-350 bp average length using a Bioruptor UCD-200 sonication, following manufacturer recommendations (Diagenode). WGS libraries were prepared using the xGen ssDNA & Low-Input DNA Library Preparation Kit and xGen UDI Primers (Integrated DNA Technologies; IDT), following kit protocol. QC of cDNA libraries was conducted using Qubit dsDNA HS assay (Life Technologies) and Bioanalyzer dsDNA HS assay (Agilent Technologies). Libraries were pooled to estimated equimolar fractions, then library QC and pooling balance were evaluated using iSeq were pooled and sequenced on Illumina iSeq. Library pools were normalized by iSeq read count, then sequenced on Illumina NovaSeqX (150 bp paired-end reads).

### DNASeq Human Detection Pipeline

Raw reads were trimmed for adapter contamination and low-quality bases using Trimmomatic (v0.39) with the *TruSeq3-PE.fa* adapter file and the parameters *ILLUMINACLIP:TruSeq3-PE.fa:2:30:10 HEADCROP:12 SLIDINGWINDOW:4:15 MINLEN:15*. As an additional filter to remove low complexity reads sequences, the trimmed reads were filtered with bbduk from the bbtools (v39.06) package. Next, the filtered and trimmed reads were mapped to the human reference genome GRCh38 with bwa-mem2 (v2.2.1), and unmapped reads were discarded in order to retain only human sequences and discard microbial sequences. As an additional validation step, all reads were compared to GRCh38 using BLAST (v2.15) with the parameters *–evalue 1e-10 –max_target_seqs 1 –max_hsps 1* and only reads with significant hits were retained. The passing reads were then remapped to GRCh38 using Minimap2 (v2.30) with the parameters -ax sr. The reads were not assembled because extracellular vesicles are expected to contain sequence fragments not whole genomes. The BAM files from Minimap2 were filtered for high quality and uniquely mapped alignments using samtools (v1.21) parameters: *view -b -q 20.* The mapping distribution of the aligned reads was visualized in R using karyoploteR (v1.36.0). Reads were quantified using the multiBigwigSummary tool from the deepTools (v3.5.6) package and providing a BED file of known genes obtained from the UCSC Table Browser (Gencode v49 knownGene). Pipeline is illustrated in **Supplemental Figure 5b**.

### DNAseq EBV Detection Pipeline

The trimmed FASTQ files from Trimmomatic (v0.39) were pushed through PathSeqPipelineSpark tool from GATK (v4.6) resulting in PathSeq scores tables. The score tables were imported into R (v4.4), and a Poisson distribution was fitted using the vcd package (v1.4) to determine the expected distribution of microbial scores. Based on this model, we considered samples with a PathSeq score of three or greater for any microbe to be a valid hit. For each sample with a PathSeq score of three or greater for EBV we extracted the reads identified by PathSeqPipelineSpark. As an additional validation step, the EBV genome (NC_007605.1) was converted into a Hidden Markov Model (HMM) profile using hmmbuild from the hmmer (v3.4) package. The HMM profile was queried against EBV candidate reads using nhmmer to verify motif-level matches. Reads identified as motif-levels matches were extracted with esl-reformat. Passing reads were then mapped to the EBV genome using Minimap2 (v2.30) with the *-ax sr* parameter. The BAM files were filtered for uniquely mapped, and high-quality alignments using samtools. EBV-mapped reads were visualized in R using karyoploteR (v1.36.0) and Gviz (v1.54). Quantification of EBV read coverage was performed with multiBigwigSummary from deepTools with a BED file of annotated genes from the EBV genome (NC_007605.1).

### Super Resolution Microscopy of EVs

Super resolution imaging of sEVs was performed using the ONi (Oxford Nanoimaging) Nanoimager and the EV Profiler 2 Kit according to a modified protocol. ONi chips were prepared according to manufacturer’s instructions using α-CD63, α-CD9, and α-CD81 tetraspanin antibodies for EV capture. For detection of EdU in EVs, sEVs generated from SLCLs grown in media containing EdU were captured on the ONi chip, fixed, and stained using the Click-it Plus EdU (ThermoFisher) reaction components containing 1x Click-iT reaction buffer, copper protectant, Alexa Fluor 488 picolyl azide, and 1x reaction buffer additive (ratios according to manufacturer’s instructions) with permeabilization buffer (ONi kit). Chips were incubated for 30 mins at room temperature, followed by washing and subsequent staining with α-CD81 (647; ONi kit) and 2x SPY555-Actin (Cytoskeleton, Inc.) for 2 hr at room temperature, protected from light. Post staining, stained EVs were washed, fixed, and subjected to dSTORM imaging using ONi NimOS software and nanoimager using 488, 560, and 640 lasers. The instrument was set to 31°C, an illumination angle of 51°, and 3000 total frames per field of view were captured.

For imaging of EBER1 ncRNA in EVs, SLCL sEVs were captured by tetraspanins on ONi chips, fixed, washed with 2x SSC/10% formamide buffer, and stained with a hybridization buffer containing 250 nM EBER1-3’ probe (CAG AGT CTG GGA AGA CAA CCA CAG ACA CCG /3AlexF647N/; Integrated DNA Technologies (IDT)), 250 nM EBER1-5’ probe (/5Alex647N/CA GAG TCT GGG AAG ACA ACC ACA GAC ACC G; IDT), hybridization solution (ready-to-use; Sigma), permeabilization buffer and staining buffer (ONi kit). EBER1 probes were designed from previous publication.^41,42^ Chips were heated to 70°C for 15 mins followed by an overnight incubation at 4°C. Next day, chips were washed and subsequently stained with Tetraspanin Trio (α-CD63, α-CD9, and α-CD81 mix; 561; ONi kit) and permeabilization buffer for 50 mins at room temperature. After staining, chips were washed, fixed, and imaged using dSTORM as described. Analysis of images was performed in CODI (ONi cloud analysis platform; https://alto.codi.bio/).

### EV RNA Isolation and RNA Sequencing

EV RNA was isolated from UC-isolated EVs using the Exosomal RNA Isolation Kit (Norgen Biotek). Heat-labile DNase was inactivated by incubation at 58°C for 5 minutes prior to library preparation. Cellular RNA was extracted from cultured cells using the Single Cell RNA Purification Kit (Norgen Biotek). On-column DNase treatment was performed during both EV and cell RNA extractions using RNase-Free DNase I Kit (Norgen Biotek). RNA quantity and quality were assessed by Qubit RNA HS (ThermoFisher) or Quant-iT RiboGreen RNA assays. RNA was stored at –80 °C until used for library preparation.

Total RNA-seq libraries were prepared using the SMARTer Stranded Total RNA-Seq Kit v3 - Pico Input Mammalian (Takara Bio), following manufacturer protocol, using 50 pg – 10 ng RNA as input. For libraries prepared from intact EVs, the kit protocol option for intact cells was followed, with 1e9 EVs used as input. QC of cDNA libraries was conducted using Qubit dsDNA HS assay (Life Technologies) and Bioanalyzer dsDNA HS assay (Agilent Technologies). Libraries were pooled to estimated equimolar fractions, then library quality and pooling balance were evaluated by sequencing on Illumina iSeq. Library pools were normalized by iSeq read count and sequenced on Illumina NovaSeqX (150 bp paired-end reads).

### RNASeq Human Detection Pipeline

In order to detect and quantify human sequences in the RNAseq extracellular vesicle data we used a similar pipeline to the *DNASeq Human Detection Pipeline*. Raw reads were filtered for low quality bases, adapters, and low complexity using Trimmomatic (v0.39) and bbduk. The filtered and trimmed reads were mapped to GRCh38 with STAR (v2.7.11b) and unmapped reads were discarded. BLAST (v2.15) was implemented as an additional validation step. Passing reads were remapped to GRCh38 using Minimap2 (v2.30) with the parameters *-ax splice:sr*. Mapped reads were filtered for quality and unique mapping with samtools (v1.21). The aligned reads were visualized in R (v4.4) with karyoploteR (v1.36.0). Transcript-level quantification was performed using Salmon (v1.10.1). Next, the transcript-level counts were imported and length-scaled normalization (countsFromAbundance = “lengthScaledTPM”) was applied using the tximport package (v1.38.2) in R to obtain gene-level abundance estimates. This normalization both within and across samples produced gene-level estimates to assess the different biotypes present which were counted with FeatureCounts from the R package Rsubread (v1.22.2). Pipeline is illustrated in **Supplemental Figure 5c.**

### RNASeq EBV Detection Pipeline

The detection and quantification of EBV sequences in the RNAseq data follows the same pipeline implemented in the *DNAseq Epstein-Barr Virus Detection Pipeline* with the following minor differences: 1) Minimap2 (v2.30) was run with the following parameter change: *-ax splice:sr*, and 2) reads were normalized within and across samples as described above with Salmon (v1.10.1) and tximport package (v1.38.2).

### Droplet Digital PCR (ddPCR)

RNA was extracted from prepared sEVs using the Norgen Exosomal RNA Isolation Kit (Norgen Biotek) according to the manufacturer’s instructions. Quantification of RNA concentration and purity (260/280) was determined using a NanoDrop 2000 Spectrophotometer (ThermoFisher). Following quantification, RNA was treated with DNase to remove residual DNA using the TURBO DNA-free kit (ThermoFisher) according to manufacturer’s instructions. DNase-treated RNA was used immediately for cDNA conversion or kept at -80°C.

DNase-treated RNA was converted to cDNA using the High-Capacity cDNA Reverse Transcription Kit (ThermoFisher) according to manufacturer’s instructions. cDNA conversion included an RT Control containing DNase-treated RNA without reverse transcription reagents. All samples were incubated on a GeneAmp PCR System 9700 (Applied Biosystems) according to the following protocol: 25°C for 10 min, 37°C for 2 hr, 85°C for 5min, 4°C hold. Following completion of the thermocycler incubation, cDNA samples were treated with RNase H and incubated at 37°C for 20 mins to remove residual RNA. RNase-treated cDNA samples were used immediately for ddPCR or stored at -20°C until use.

For ddPCR, RNase-treated cDNA samples were combined with EBV target (*EBER1*, *BZLF1*/Zta, or *BFRF3*) 20x primer/probe mix (FAM), HPRT 20x primer/probe mix (VIC), ddPCR SuperMix (BioRad), and PCR water (Quality Biological). All samples were run in triplicate. Droplets were generated using a QX200 Droplet Generator (BioRad), with approximately 20,000 droplets generated per well. Droplets were incubated on a GeneAmp PCR System 9700 Thermocycler for amplification of target genes: 95°C for 10 min, 94°C for 30 sec, and 59°C for 1 min x 40 cycles, followed by 98°C for 10 min, 12°C hold. Following completion of the PCR amplification cycle, gene expression was analyzed using a QX200 Droplet Reader (BioRad) and QuantaSoft software version 1.7.4.0917 (BioRad). All primers and probes were obtained from ThermoFisher. Probes were Custom TaqMan probes and primers were Unlabeled Sequence Detection Primers unless otherwise indicated in **Table 1**. Sequences for primers and probes are provided in **Table 1**. EBER1 primers and probes were designed using Primer3 and validated for specificity. Copy numbers of EBV targets were normalized to EV input numbers as determined by MRPS.

**Table 1:**
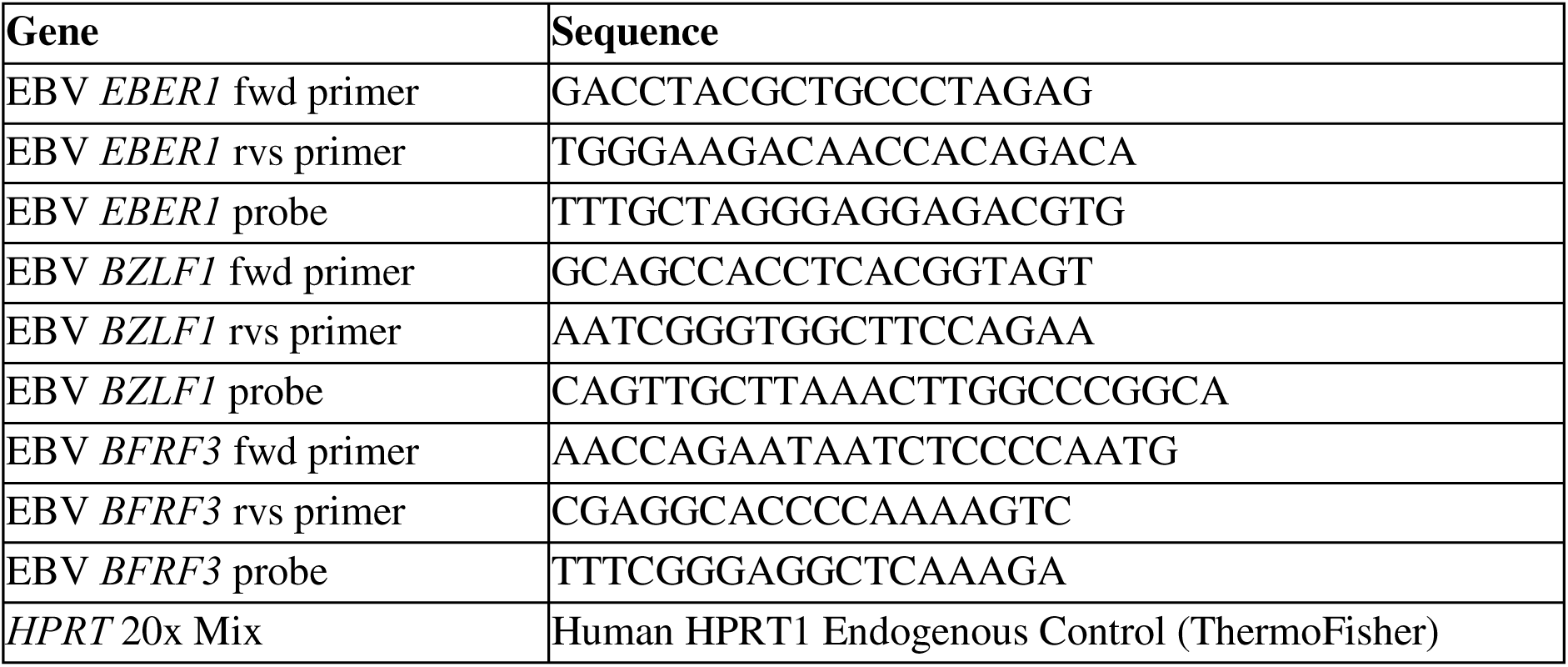
Primer and probe sequences for ddPCR application.

### Use of AI-Assisted Tools

Custom R scripts were developed with assistance from Claude Sonnet 4.5 to 4.7 (Anthropic), and generative AI using HHS Enterprise ChatGPT (OpenAI) and Claude was used in a limited capacity to refine manuscript text for clarity and style. All AI-assisted outputs were reviewed and verified by the authors, who executed the analyses and take full responsibility for the manuscript.

## Author Contributions

MLP, KvK-J, KJ, JCJ, and SJ conceptualized and designed the work. MLP, SC, EHS, JG, CVD, AD, JP, EH, TKM, WY, and R-CH acquired the data. MLP, RP, SC, EHS, MRD, CVD, JP, EH, WY, LMJ, and R-CH analysed the data. MLP, RP, SC, EHS, MRD, CVD, JP, EH, C-SH, LMJ, R-CH, ICG, KvK-J, KJ, JCJ, and SJ interpreted the data. MLP, RP, SC, JP, EH, ICG, KvK-J, KJ, and JCJ created and/or implemented analysis pipelines. MLP, RP, SC, MRD, JP, EH, LMJ, JCJ, and SJ wrote the manuscript. MLP, CVD, C-SH, ICG, KJ, JCJ and SJ edited the manuscript. NN, JO, JS, and SJ acquired, stored, and organized samples. MCM generated cell lines and reagents used in this study.

## Acknowledgments

We would like to acknowledge and thank all patients, volunteers, and clinical staff at the Neuroimmunology Clinic who contributed to this project.

