## Supplemental Information for "Extracellular vesicles from wild-type Epstein–Barr virus–transformed B-cells export host DNA and EBV EBER1"

**Supplemental Table 1: EBV proteins detected in SLCL-derived sEVs by DIA mass spectrometry.**

Table listing EBV-encoded proteins and their associated descriptive names (Protein Description) identified in sEV preparations by DIA mass spectrometry across individual SLCL samples with corresponding abundance values. Positive (Namalwa) and negative (Ramos) control B-cell lines were included. Undetected proteins are denoted by " - ".

| Protein | Protein Description | ND2 | ND3 | ND5 | SMS2 | SMS7 | AMS2 | AMS3 | AMS4 | Namalwa | Ramos |
|---|---|---|---|---|---|---|---|---|---|---|---|
| **MCP** | Major capsid protein | 1.35E+04 | 4.15E+04 | 9.06E+04 | 3.04E+05 | 9.11E+04 | 2.37E+04 | 9.52E+04 | 1.48E+05 | 2.38E+04 | - |
| **BLRF2** | Tegument protein BLRF2 | - | - | 5.49E+04 | 1.74E+05 | 1.32E+05 | 4.64E+04 | 5.12E+04 | 1.20E+05 | - | - |
| **BBRF1** | Portal protein | - | - | 2.27E+04 | 5.00E+04 | 2.08E+04 | - | 2.92E+04 | 1.01E+05 | - | - |
| **BVRF2/BdRF1** | Capsid scaffolding protein | - | - | 6.23E+04 | 1.81E+05 | 4.09E+04 | 1.52E+04 | 5.35E+04 | 9.63E+04 | - | - |
| **BMRF1** | DNA polymerase processivity factor BMRF1 | 1.02E+04 | - | 4.88E+04 | 8.29E+04 | 4.78E+04 | 1.03E+04 | 6.43E+04 | 6.30E+04 | - | - |
| **TRX2** | Triplex capsid protein 2 | - | - | 2.83E+04 | 9.88E+04 | 2.08E+04 | 4.28E+03 | 2.58E+04 | 4.67E+04 | - | - |
| **RIR2** | Ribonucleoside-diphosphate reductase small subunit | 1.23E+04 | - | 5.70E+04 | 3.31E+04 | 4.22E+04 | 1.34E+04 | 5.89E+04 | 4.48E+04 | - | - |
| **BALF5** | DNA polymerase catalytic subunit | - | - | 2.86E+04 | 3.70E+04 | - | - | 2.55E+04 | 3.95E+04 | - | - |
| **gH** | Envelope glycoprotein H | - | - | 3.57E+04 | 4.23E+04 | 2.95E+04 | - | 3.00E+04 | 3.83E+04 | - | - |
| **RIR1** | Ribonucleoside-diphosphate reductase large subunit | 1.77E+04 | - | 5.43E+04 | 4.36E+04 | 3.05E+04 | 1.81E+04 | 4.96E+04 | 3.75E+04 | - | - |
| **BDLF3** | Glycoprotein BDLF3 | - | - | 1.82E+04 | 5.99E+04 | 2.02E+04 | - | 1.98E+04 | 3.11E+04 | - | - |
| **gM** | Envelope glycoprotein M | - | - | 5.04E+03 | 3.41E+04 | 7.04E+03 | - | 1.81E+04 | 2.92E+04 | 8.55E+02 | - |
| **BGLF4** | Serine/threonine-protein kinase BGLF4 | - | - | 1.89E+04 | 2.54E+04 | 2.52E+04 | - | 2.06E+04 | 2.28E+04 | - | - |
| **dUTPase** | Deoxyuridine 5'-triphosphate nucleotidohydrolase | 5.06E+03 | - | 1.84E+04 | 2.01E+04 | 2.35E+04 | - | 2.95E+04 | 2.21E+04 | - | - |
| **TRX1** | Triplex capsid protein 1 | - | - | 7.97E+03 | 3.77E+04 | 9.30E+03 | - | 1.09E+04 | 2.12E+04 | - | - |
| **SCP** | Small capsomere-interacting protein | - | - | - | 4.59E+04 | - | - | 2.42E+04 | 1.60E+04 | - | - |
| **BMRF2** | Protein BMRF2 | - | - | 1.27E+04 | 3.25E+04 | - | - | - | 1.47E+04 | - | - |
| **UNG** | Uracil-DNA glycosylase | - | - | 4.49E+04 | - | - | - | 3.36E+04 | 1.38E+04 | - | - |
| **LMP2** | Latent membrane protein 2 | 1.68E+04 | 1.52E+04 | 1.40E+04 | 1.98E+04 | 9.04E+03 | 1.80E+04 | 1.60E+04 | 1.38E+04 | - | - |
| **BNRF1** | Major tegument protein | - | - | 6.51E+03 | 2.60E+04 | 8.10E+03 | - | 1.21E+04 | 1.21E+04 | - | - |
| **EBNA-LP** | Epstein-Barr nuclear antigen leader protein | - | - | 1.29E+04 | 4.18E+03 | 1.43E+04 | 2.27E+04 | 3.93E+04 | 1.21E+04 | - | - |
| **LF2** | Protein LF2 | - | - | - | 1.28E+04 | - | - | 9.19E+03 | 1.10E+04 | - | - |
| **gB** | Envelope glycoprotein B | - | - | 5.18E+03 | 2.55E+04 | 4.63E+03 | - | 9.28E+03 | 1.08E+04 | - | - |
| **EBNA1** | Epstein-Barr nuclear antigen 1 | 3.13E+03 | 6.17E+03 | 2.40E+03 | 5.53E+03 | - | 2.66E+03 | 4.94E+03 | 8.97E+03 | - | - |
| **BDLF2** | Protein BDLF2 | - | - | 4.21E+03 | 1.34E+04 | 5.60E+03 | - | 8.72E+03 | 7.41E+03 | - | - |
| **BNLF2b** | Uncharacterized protein BNLF2b | - | - | 3.30E+03 | 7.66E+03 | 3.76E+03 | - | 3.67E+03 | 6.89E+03 | - | - |
| **BMLF1** | mRNA export factor ICP27 homolog | 5.99E+02 | - | 1.17E+04 | 2.93E+03 | 5.69E+03 | - | 1.03E+04 | 6.55E+03 | - | - |
| **BGLF5** | Shutoff alkaline exonuclease | 1.56E+03 | - | 1.44E+04 | 6.89E+03 | 8.11E+03 | - | 1.03E+04 | 5.55E+03 | - | - |
| **BARF1** | Secreted protein BARF1 | - | - | 9.40E+03 | 1.16E+04 | 5.41E+03 | - | 1.13E+04 | 5.35E+03 | - | - |
| **gH** | Envelope glycoprotein H | - | - | - | - | - | - | - | 3.79E+03 | - | - |
| **BRRF1** | Transcriptional activator BRRF1 | - | - | 1.38E+04 | 3.29E+03 | 5.11E+03 | - | 4.52E+03 | 3.58E+03 | - | - |
| **TK** | Thymidine kinase | - | - | 4.13E+03 | - | - | - | 4.20E+03 | 2.78E+03 | - | - |
| **BHRF1** | Apoptosis regulator BHRF1 | - | - | 1.72E+03 | 1.74E+03 | 9.99E+02 | - | 1.49E+03 | 2.71E+03 | - | - |
| **gL** | Envelope glycoprotein L | - | - | 1.22E+03 | 3.42E+03 | - | - | 5.52E+03 | 2.51E+03 | - | - |
| **BRRF2** | Tegument protein BRRF2 | - | - | - | 1.75E+03 | - | - | 1.88E+03 | 1.54E+03 | - | - |
| **BRLF1** | Replication and transcription activator | - | - | 1.43E+03 | - | - | - | 1.38E+03 | 1.32E+03 | - | - |
| **BZLF2** | Glycoprotein 42 | - | - | 2.22E+03 | 1.80E+03 | 1.88E+03 | - | 2.04E+03 | 9.57E+02 | - | - |
| **BALF1** | Apoptosis regulator BALF1 | - | - | - | 4.23E+03 | - | - | - | - | - | - |
| **BFLF1** | Packaging protein UL32 homolog | - | - | 5.19E+03 | 1.08E+03 | - | - | 3.95E+03 | - | - | - |
| **CVC2** | Capsid vertex component 2 | - | - | - | 1.94E+04 | - | - | - | - | - | - |

**Supplemental Table 2: EBV RNA counts detected in SLCL-derived sEVs by total stranded RNA sequencing.**

Table listing EBV-encoded transcripts detected from SLCL sEV RNASeq, with counts for each transcript across individual SLCL samples and comparator lines where indicated. EBER1 was the most abundant EBV transcript across sEV preparations. EBV positive control cell line Namalwa was included. Undetected RNAs are denoted by " - ".

| Product | Namalwa | ND2 | ND3 | ND5 | SMS2 | SMS7 | AMS2 | AMS3 | AMS4 |
|---|---|---|---|---|---|---|---|---|---|
| EBER-1 | 234 | 1489 | 3197 | 1960 | 1260 | 3788 | 4049 | 4777 | 276 |
| EBNA-3B | 218 | 33 | 167 | 1171 | 447 | 836 | 398 | 1139 | 584 |
| LMP-1 | - | 6 | 46 | 1947 | 423 | 441 | 451 | 845 | 520 |
| RPMS1 | - | 184 | 62 | 896 | 513 | 510 | 182 | 1146 | 457 |
| EBNA-3C | 218 | 23 | 144 | 854 | 266 | 650 | 377 | 910 | 436 |
| EBNA-2 | 294 | 76 | 138 | 716 | 515 | 729 | 258 | 810 | 267 |
| EBNA-3A | 218 | 24 | 129 | 753 | 364 | 454 | 237 | 885 | 486 |
| EBNA-1.1 | 218 | 21 | 130 | 504 | 90 | 475 | 179 | 605 | 263 |
| A73 | - | 134 | 34 | 530 | 394 | 196 | 99 | 642 | 230 |
| ebv-miR-BHRF1-3 | - | 62 | 45 | 653 | 456 | 251 | 27 | 486 | 168 |
| EBNA-LP | 211 | 53 | 80 | 434 | 124 | 413 | 185 | 599 | 111 |
| Rta | 1 | 17 | 6 | 489 | 177 | 373 | 82 | 403 | 322 |
| EBER-2 | 17 | 55 | 307 | 275 | 127 | 221 | 117 | 284 | 50 |
| LMP-2B | 15 | 8 | 82 | 189 | 120 | 108 | 71 | 162 | 347 |
| EBNA-1 | 9 | - | 8 | 273 | 97 | 173 | 24 | 232 | 93 |
| Zta | - | - | - | 265 | 69 | 91 | 77 | 204 | 106 |
| ebv-miR-BHRF1-2 | - | 12 | - | 302 | 167 | 54 | 24 | 172 | 47 |
| ebv-miR-BHRF1-2* | - | 12 | - | 166 | 143 | 60 | 24 | 121 | 54 |
| LMP-2A | - | 2 | - | 102 | 27 | 53 | - | 150 | 110 |
| ebv-miR-BART17-5p | - | - | - | - | 51 | 16 | - | 90 | 22 |
| ebv-miR-BART15 | - | - | - | - | 8 | 10 | 23 | 98 | 35 |
| ebv-miR-BART6-5p | - | - | - | - | 35 | 27 | - | 91 | 10 |
| ebv-miR-BART6-3p | - | - | 1 | 1 | 5 | 30 | - | 84 | 17 |
| ebv-miR-BART17-3p | - | - | - | - | 35 | 16 | - | 71 | 14 |
| ebv-miR-BART16 | - | 9 | - | - | 2 | 7 | - | 78 | 19 |
| ebv-miR-BART14* | - | - | - | 38 | 5 | 14 | - | 16 | 23 |
| ebv-miR-BART5* | - | - | - | - | 5 | - | - | 55 | 15 |
| RPMS1-25 | - | - | - | 5 | 11 | 1 | - | 15 | 31 |
| ebv-miR-BHRF1-1 | - | - | - | 20 | 1 | 7 | - | 23 | 9 |
| ebv-miR-BART5 | - | - | - | - | 2 | - | - | 39 | 10 |
| ebv-miR-BART13* | - | - | 4 | 3 | - | 23 | - | 11 | 8 |
| ebv-miR-BART9 | - | - | - | - | - | - | - | - | 43 |
| ebv-miR-BART9* | - | - | - | 1 | - | - | - | - | 41 |
| ebv-miR-BART8 | - | - | - | 1 | - | 1 | - | 14 | 22 |
| ebv-miR-BART13 | - | - | - | - | - | 20 | - | 11 | 4 |
| ebv-miR-BART19-5p | - | - | - | - | - | - | - | 12 | 19 |
| ebv-miR-BART4* | - | - | 2 | 1 | 10 | - | 10 | - | 8 |
| ebv-miR-BART3 | - | - | 3 | 2 | - | - | 10 | 8 | 5 |
| ebv-miR-BART10* | - | - | - | 1 | 1 | 5 | - | - | 18 |
| ebv-miR-BART8* | - | - | - | - | - | - | 2 | - | 23 |
| ebv-miR-BART18-3p | - | - | - | 18 | - | - | 1 | 2 | 1 |
| ebv-miR-BART12 | - | - | - | 18 | - | - | - | 2 | 1 |
| ebv-miR-BART11-5p | - | - | - | - | - | - | - | - | 20 |
| ebv-miR-BART10 | - | - | - | - | - | - | - | - | 19 |
| ebv-miR-BART14 | - | - | - | - | - | - | - | 2 | 17 |
| ebv-miR-BART11-3p | - | - | - | - | - | - | - | - | 18 |
| ebv-miR-BART21-5p | - | - | - | - | 1 | 3 | 2 | 2 | 9 |
| ebv-miR-BART4 | - | - | - | - | 10 | - | - | 1 | 6 |
| ebv-miR-BART19-3p | - | - | - | - | - | - | - | - | 15 |
| ebv-miR-BART2-5p | - | - | - | - | 1 | 1 | - | - | 11 |
| ebv-miR-BART20-3p | - | - | - | 4 | 2 | - | - | - | 5 |
| ebv-miR-BART2-3p | - | - | - | - | - | - | 1 | - | 10 |
| ebv-miR-BART1-3p | - | - | - | 1 | 2 | - | - | 1 | 5 |
| ebv-miR-BART1-5p | - | - | - | 1 | - | - | - | 2 | 3 |
| ebv-miR-BART20-5p | - | - | - | - | - | - | - | 1 | 5 |
| ebv-miR-BART7* | - | - | - | - | - | - | - | - | 6 |
| ebv-miR-BART3* | - | - | 3 | - | - | - | - | - | - |
| ebv-miR-BART21-3p | - | - | - | - | - | - | - | - | 2 |
| ebv-miR-BART7 | - | - | - | - | - | - | - | - | 1 |

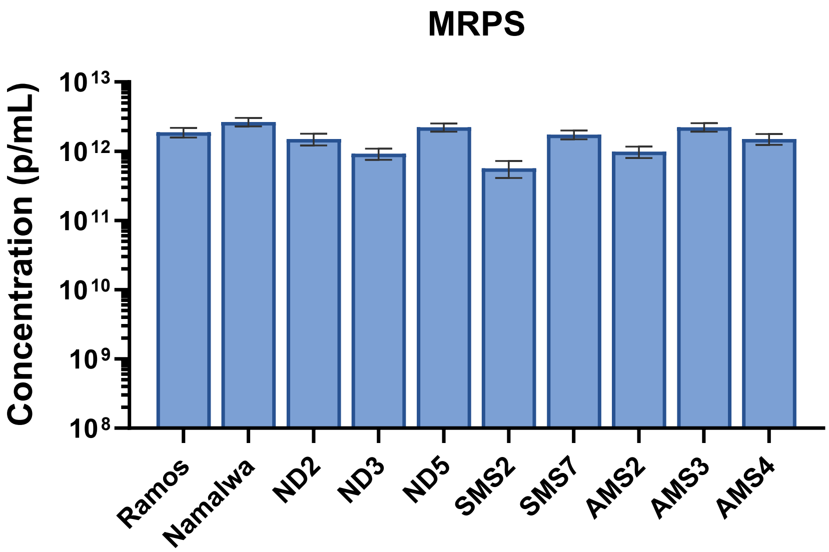


**Supplemental Figure 1: MRPS quantification of sEV preparations from SLCLs**

Microfluidic resistive pulse sensing (MRPS) quantification of sEV concentration in particles/mL (p/mL) generated from normal donor (ND), stable MS (SMS), and active MS (AMS) SLCLs. Bars represent mean particle concentration.

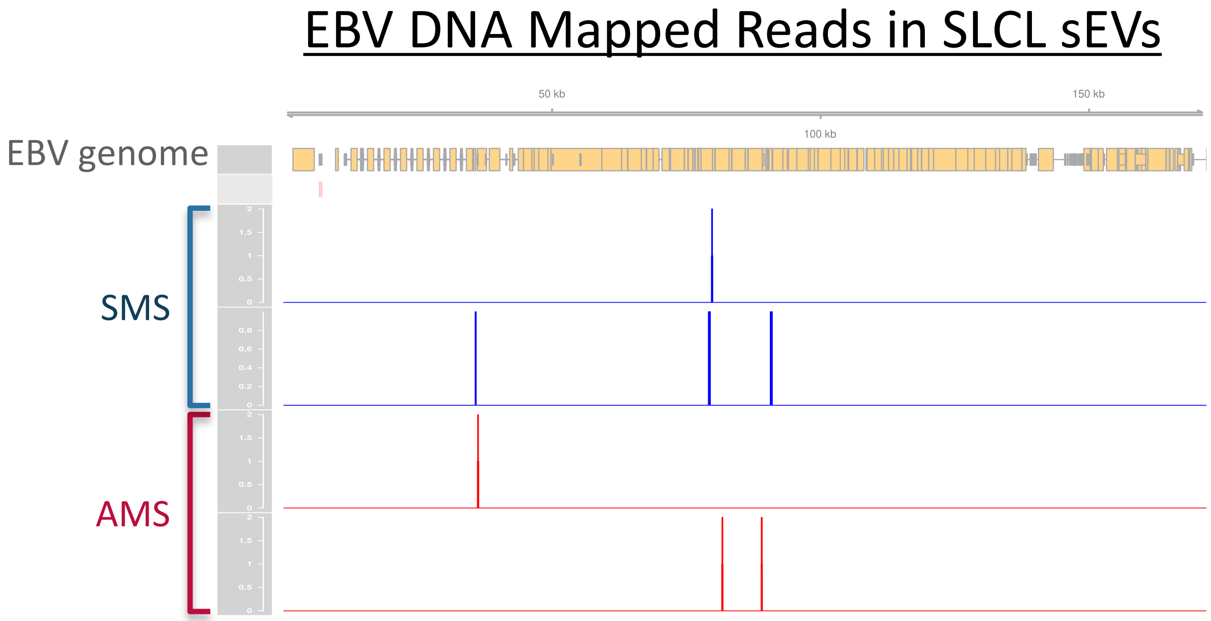


**Supplemental Figure 2: EBV genomic DNA is minimally represented in SLCL-derived sEVs.** Integrated Genome Viewer (IGV) browser view of DNASeq reads mapped to the EBV genome from SLCL-derived sEV preparations. Across few SMS and AMS samples, only sparse and discontinuous EBV-aligned reads were detected, with no consistent enrichment across specific viral genomic regions.

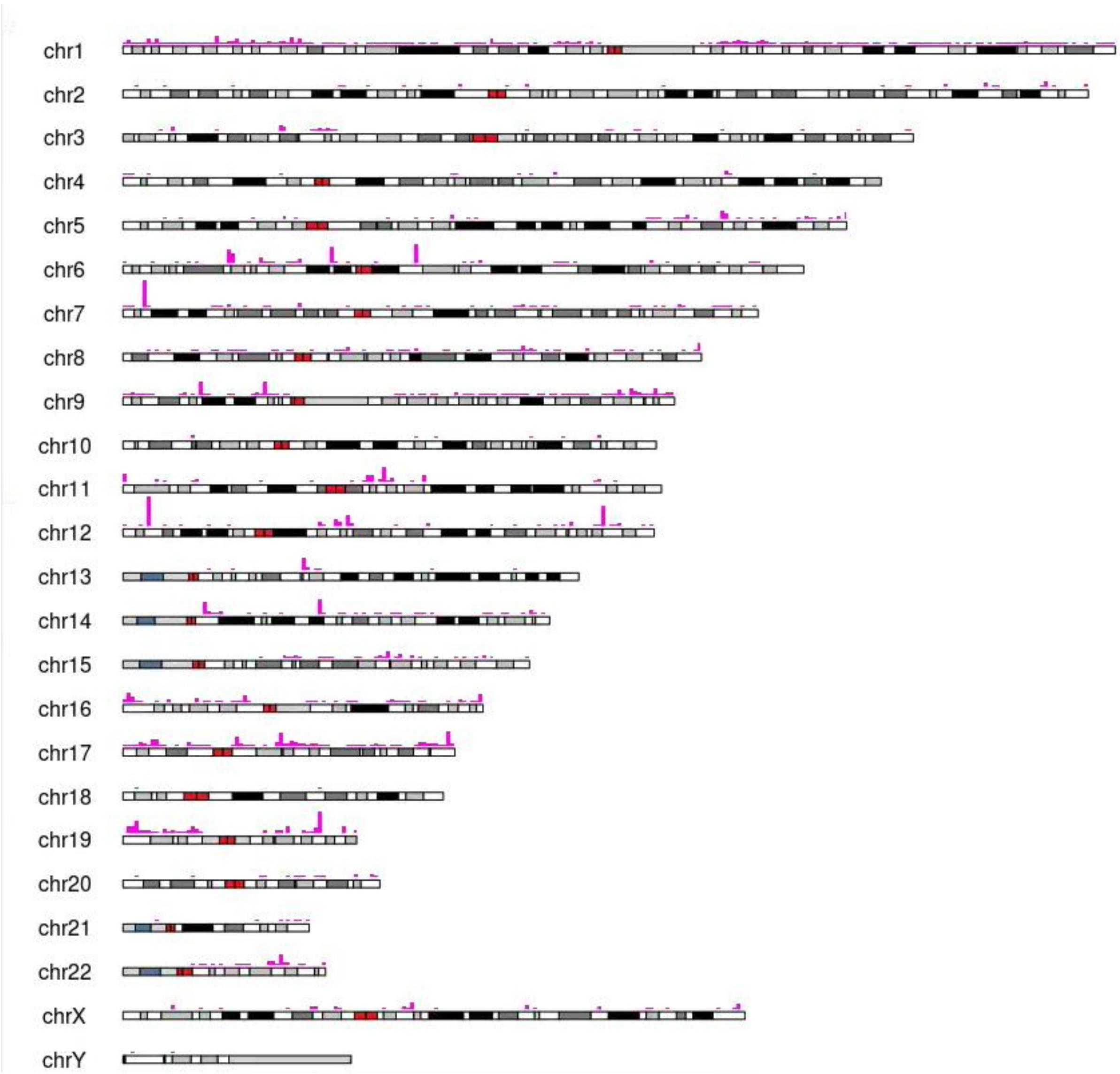


**Supplemental Figure 3: Representative chromosomal mapping of host RNA reads in SLCL-derived sEVs.**

Representative karyotype density plot showing genome-wide distribution of host-derived SLCL sEV RNASeq reads (magenta vertical bars) across human chromosomes. Reads are distributed broadly throughout the genome without a dominant positional bias. Red blocks show centromeric regions of each chromosome. Black and light-gray blocks represent the chromosome ideogram/cytoband structure.

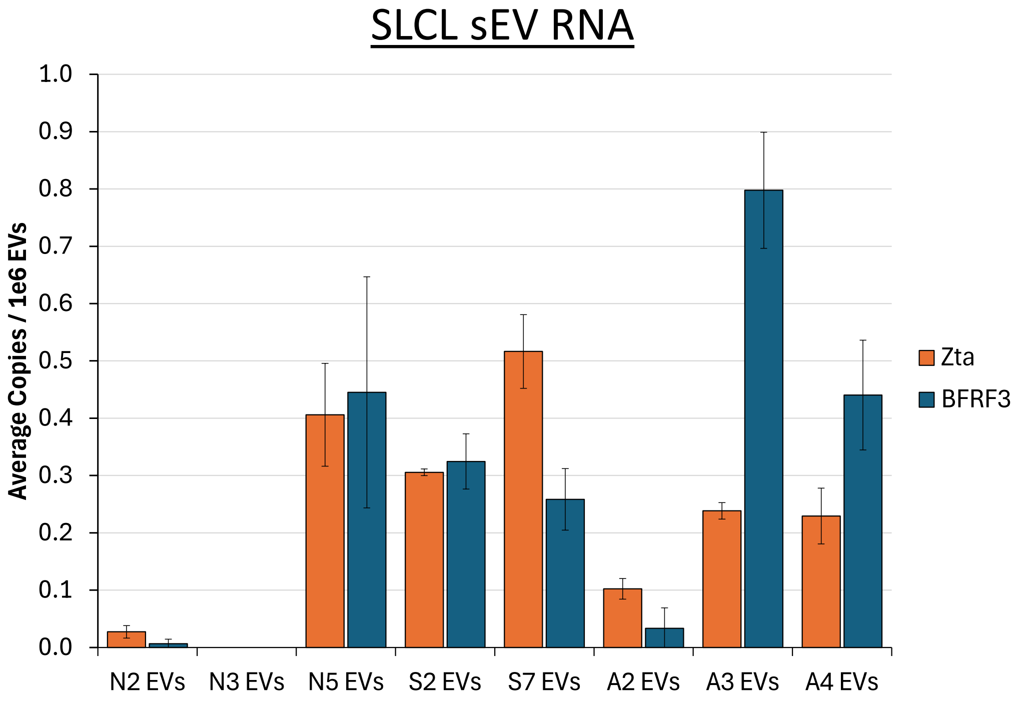


**Supplemental Figure 4: Detection of low-abundance lytic EBV transcripts in SLCL-derived sEV RNA.**

Droplet digital PCR quantification of the lytic EBV transcripts BZLF1 (Zta; orange) and BFRF3 (blue) from sEV RNA from individual SLCL lines, reported as average copies per $10^6$ EVs.

**A)** DIA Proteomics sEV Analysis Pipeline

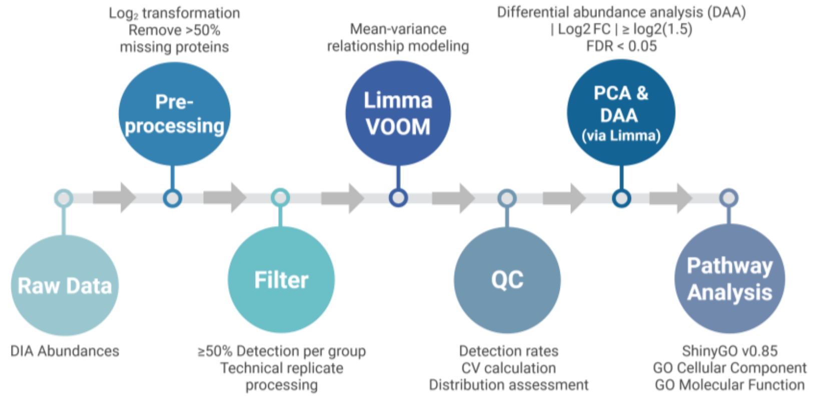


**B)** WGS DNASeq Human sEV Analysis Pipeline

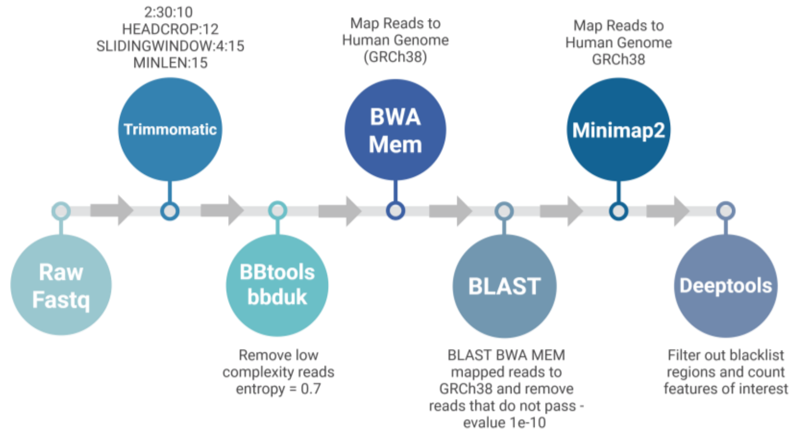


**C)** Total Stranded RNASeq Human sEV Analysis Pipeline

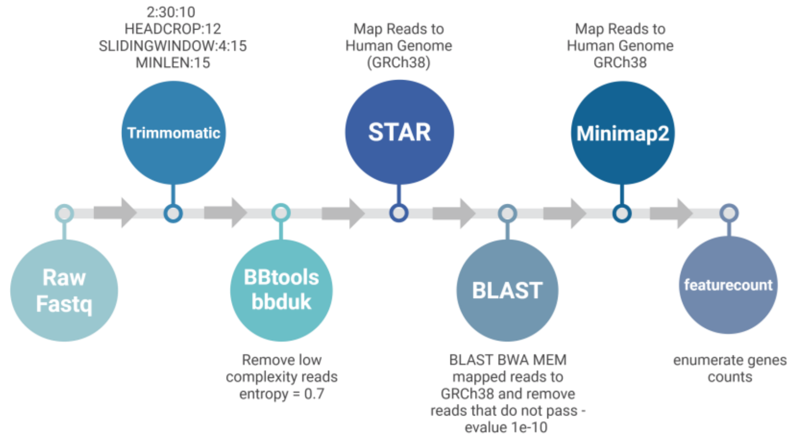

**Supplemental Figure 5: Bioinformatic pipelines used for multiomic analysis of SLCL-derived sEV cargo.**

**A)** Schematic of the DIA proteomics analysis workflow, including preprocessing, filtering, normalization with limma/voom, quality control, differential expression analysis, and pathway enrichment analysis. **B)** Schematic of the whole-genome sequencing human DNA analysis pipeline, including adapter and quality trimming, low-complexity filtering, alignment to GRCh38, BLAST-based validation, remapping, and downstream quantification/visualization. **C)** Schematic of the total stranded RNASeq human analysis pipeline, including read trimming, low-complexity filtering, alignment to GRCh38 with STAR, BLAST-based validation, remapping, and downstream quantification/visualization. Pipeline figures were created using BioRender.
